# Homocysteine Metabolites Inhibit Autophagy and Elevate Amyloid Beta by Impairing Phf8/H4K20me1-dependent Epigenetic Regulation of mTOR in Cystathionine β-Synthase-Deficient Mice

**DOI:** 10.1101/2023.03.22.533769

**Authors:** Łukasz Witucki, Hieronim Jakubowski

**Affiliations:** Department of Microbiology, Biochemistry and Molecular Genetics, Rutgers University, New Jersey Medical School, Newark, NJ 07103, USA; Department of Biochemistry and Biotechnology, Poznań University of Life Sciences, Poznań, Poland

**Keywords:** APP, amyloid beta, *Cbs*^-/-^ mouse, homocysteine thiolactone, *N*-homocysteinylated protein, Phf8, H4K20me1, mTOR signaling, autophagy

## Abstract

The loss of cystathionine β-synthase (CBS), an important homocysteine (Hcy)-metabolizing enzyme or the loss of PHF8, an important histone demethylase participating in epigenetic regulation, causes severe mental retardation in humans. Similar neuropathies were also observed in *Cbs*^-/-^ and *Phf8*^-/-^ mice. How CBS or PHF8 depletion can cause neuropathy was unknown. To answer this question, we examined a possible interaction between PHF8 and CBS using *Cbs*^-/-^ mouse and neuroblastoma cell models. We quantified gene expression by RT-qPCR and Western blotting, mTOR-bound H4K20me1 by chromatin immunoprecipitation (CHIP) assay, and amyloid β (Aβ) by confocal fluorescence microscopy using anti-Aβ antibody. We found significantly reduced expression of Phf8, increased H4K20me1, increased mTOR expression and phosphorylation, and increased App, both on protein and mRNA levels in brains of *Cbs*^-/-^ mice *vs. Cbs*^+/-^ sibling controls. Autophagy-related proteins Becn1, Atg5, and Atg7 were downregulated while p62 was upregulated on protein and mRNA levels, suggesting impaired autophagy in *Cbs*^-/-^ brains. In mouse neuroblastoma N2a or N2a-APPswe cells, treatments with Hcy-thiolactone, *N*-Hcy-protein or Hcy, or *Cbs* gene silencing by RNA interference significantly reduced Phf8 expression and increased total H4K20me1 as well as mTOR promoter-bound H4K20me1. This led to transcriptional mTOR upregulation, autophagy downregulation, and significantly increased App and Aβ levels. The *Phf8* gene silencing increased Aβ, but not App, levels. Taken together, our findings identify Phf8 as a regulator of Aβ synthesis and suggest that neuropathy of Cbs deficiency is mediated by Hcy metabolites, which transcriptionally dysregulate the Phf8->H4K20me1->mTOR->autophagy pathway thereby increasing Aβ accumulation.

## 1 INTRODUCTION

The sulfur-containing amino acids methionine (Met), homocysteine (Hcy), and cysteine (Cys) are metabolically related and play important roles in cellular physiology (1). Met is an essential amino acid that participates in the biosynthesis of proteins (>20,000) and *S*-adenosylmethionine (AdoMet). AdoMet provides methyl groups for the biological methylation reactions (>300) and propyl groups for the polyamine spermidine and spermine biosynthesis. The methylation reactions generate *S*-adenosylhomocysteine (AdoHcy). Enzymatic hydrolysis of AdoHcy is the only known metabolic source of Hcy. Re-methylation of Hcy, which regenerates Met, is essential for the folate and one-carbon metabolism, which provides one-carbon units for nucleotides required for DNA and RNA biosynthesis. Transsulfuration of Hcy generates Cys, a semi-essential amino acid, participating in the biosynthesis of proteins, glutathione, hydrogen sulfide, and taurine. Hcy is also metabolized by methionyl-tRNA synthetase to Hcy-thiolactone (2-5), which modifies proteins in a nonenzymatic *N*-homocysteinylation reaction that generates *N*-Hcy-protein (5, 6).

Elevated levels of Hcy due to nutritional or genetic deficiencies are associated with neuropathology affecting the central nervous system (CNS). Cystathionine β-synthase (CBS) deficiency due to mutations in the *CBS* gene is the most prevalent inborn error in the sulfur amino acid metabolism that results in severe hyperomocysteinemia (HHcy) characterized by elevated levels of Hcy and its metabolites such as Hcy-thiolactone and *N*-Hcy-protein (1). CBS deficiency affects the CNS and causes severe mental retardation, learning disability, reduced IQ (7), psychosis, obsessive-compulsive and behavior/personality disorders (8). Accelerated brain atrophy related to elevated plasma Hcy has been reported in patients with Alzheimer’s disease (9), in patients with alcoholism (10), and in healthy elderly individuals (11). These phenotypes were replicated in *Cbs*^-/-^ mice on the C57BL/6J background, which show cognitive impairment manifested as reduced problem-solving abilities, learning disability, and short-term and long-term memory in the puzzle box test (12). *Cbs*^-/-^ mice on the C3H/HeJ background show cerebellar, although not cerebral, malformation and a transient learning deficit on day 2 in the passive avoidance step-through test (13).

Molecular bases of neurological impairments in CBS deficiency are not known. Although Hcy, Hcy-thiolactone, and *N*-Hcy-proteins accumulate in CBS-deficient patients and mice (14-17), it is not known how each of these metabolites can contribute to neuropathy associated with CBS deficiency.

Individuals with elevated plasma total Hcy show accelerated brain atrophy, impaired cognition, and are at higher risk and developing Alzheimer’s disease (18). Alzheimer’s disease patients show upregulated brain mTOR signaling (19).

Plant homeodomain finger protein 8 (PHF8) has been identified as one of the X chromosome genes linked to intellectual disability syndrome, autism spectrum disorder, attention deficit hyperactivity disorder (20), and severe mental retardation (21). PHF8 is a histone demethylase that can demethylate H4K20me1, H3K9me2/me1, andH3K27me2. Demethylation of H4K20me1 by PHF8 is important for maintaining homeostasis of mTOR signaling. The human PHF8 deficiency phenotype has been replicated in *Phf8*^-/-^ mice, which show impaired hippocampal long-term potentiation and behavioral deficits in learning and memory (22).

How CBS or PHF8 depletion can cause neuropathy was unknown. The present work was undertaken to test a hypothesis that CBS depletion reduces expression of PHF8, increases H4K20me1, mTOR expression and phosphorylation, inhibits autophagy, and increases amyloid β (Aβ) accumulation. Towards this end, we examined Phf8 expression and H4K20me1 levels in *Cbs*^-/-^ mice. We also used mouse neuroblastoma N2a and Aβ-overproducing N2a-APPswe cells to study how individual Hcy metabolites or depletion of Cbs or Phf8 by RNA interference affect Phf8 expression and its downstream effects on expression of mTOR and autophagy-related proteins, as well as App expression and Aβ accumulation.

## 2 MATERIALS AND METHODS

### 2.1 Mice

Transgenic *Tg-I278T Cbs*^-/-^ mice (23, 24) on the C57BL/6J background were housed and bred at the Rutgers-New Jersey Medical School Animal Facility. Transgene harboring human *CBS I278T* (*Tg-I278T*) variant was used to rescue the neonatal lethality associated with homozygosity for *Cbs*^-/-^. In these animals, the human mutant CBS, which has less than 3% of the activity of wild-type enzyme, is under control of the zinc-inducible metallothionein promoter, which allows to rescue the neonatal lethality phenotype of *Cbs*^-/-^ in mice by supplementing the drinking water of pregnant dams with 25 mM zinc chloride. Zinc-water was replaced by plain water after weaning at 30 days. Only *Cbs*^-/-^ mice exhibit changed phenotype characterized by facial alopecia, thin, smooth, shiny tail, reduced body weight and shortened life span compared to *Cbs*^+/-^ animals (23, 24). Mouse *Cbs* genotypes were established by PCR using the following primers: forward 5′-GGTCTGGAATTCACTATGTAGC-3′, wild type reverse 5′-CGGATGACCTGCATTCATCT*-*3′, mutant reverse: 5′-GAGGTCGACGGTATCGATA-3′. The mice were fed with a standard rodent chow (LabDiet5010; Purina Mills International, St. Louis MO, USA).

Two- to 12-month-old *Cbs*^-/-^ mice were used in experiments. Control animals were *Cbs*^+/-^ siblings. Animal procedures were approved by the Institutional Animal Care and Use Committee at Rutgers-New Jersey Medical School.

### 2.2 Brain protein extraction

Mice were euthanized by CO_2_ inhalation, the brains collected and frozen on dry ice. Frozen brains were pulverized with dry ice using a mortar and pestle and stored at −80°C. Proteins were extracted from the pulverized brains (50±5 mg) using RIPA buffer (4 v/w, containing protease and phosphatase inhibitors) with sonication (Bandelin SONOPLUS HD 2070) on wet ice (three sets of five 1-s strokes with 1 min cooling interval between strokes). Brain extracts were clarified by centrifugation (15,000 g, 15 min, 4°C) and clear supernatants containing 8-12 mg protein/mL were collected. Protein concentrations were measured with BCA kit (Thermo Scientific).

### 2.3 Cell culture and treatments

Mouse neuroblastoma N2a and N2a-APPswe cells, harboring a human APP transgene with the K670N and M671L Swedish mutations (25) were grown in DMEM/F12 medium (Thermo Scientific) supplemented with 5% FBS, non-essential amino acids, and antibiotics (MilliporeSigma) (37°C, 5% CO_2_).

After reaching 70-80% confluency, cell monolayers were washed 2-times with PBS and overlaid with DMEM medium without Met (Thermo Scientific), supplemented with 5% dialyzed FBS (MilliporeSigma) and non-essential amino acids. D,L-Hcy, L-Hcy-thiolactone (MilliporeSigma), or *N*-Hcy-protein (prepared by modification of FBS proteins with Hcy-thiolactone as described in ref. (26)) were added (at concentrations indicated in figure legends) and the cultures were incubated at 37°C in 5% CO_2_ atmosphere for 24 h.

For gene silencing, siRNAs targeting the *Cbs* (Cat. # 100821 and s63474) and Phf8 (Cat. # S115808, and S115809) (Thermo Scientific) were transfected into cells maintained in Opti-MEM medium by 24-h Lipofectamine RNAiMax (Thermo Scientific) treatments. Cellular RNA for RT-q PCR analysis were isolated as described in section 2.5 below. For protein extraction, RIPA buffer (MilliporeSigma) was used according to manufacturer’s protocol.

### 2.4 Western blots

Proteins were separated by SDS-PAGE on 10% gels (20 μg protein/lane) and transferred to PVDF membrane (Bio-Rad) for 20 min at 0.1 A, 25 V using Trans Blot Turbo Transfer System (Bio-Rad). After blocking with 5 % bovine serum albumin in TBST buffer (overnight, room temperature), the membranes were and incubated with anti-Phf8 (Abcam, ab36068), anti-H4K20me1 (Abcam ab177188), anti-mTOR (CS #2983), anti-pmTOR Ser2448 (CS, #5536), anti-Atg5 (CS, #12994), anti-Atg7 (CS, #8558), anti-

Beclin-1 (CS, #3495), anti-p62 (CS, #23214), anti-Gapdh (CS, #5174), anti-Cbs (Abcam, ab135626), or anti-App (Abcam, ab126732) or for 1 hour. Membranes were washed three times with TBST buffer, 10 min each, and incubated with goat anti-rabbit IgG secondary antibody conjugated with horseradish peroxidase. Positive signals were detected using Western Bright Quantum-Advansta K12042-D20 and GeneGnome XRQ NPC chemiluminescence detection system. Bands intensity was calculated using Gene Tools program from Syngene.

### 2.5 RNA isolation, cDNA synthesis, RT-qPCR analysis

Total RNA was isolated using Trizol reagent (MilliporeSigma). cDNA synthesis was conducted using Revert Aid First cDNA Synthesis Kit (Thermo Fisher Scientific) according to manufacturer’s protocol. Nucleic acid concentration was measured using NanoDrop (Thermo Fisher Scientific). RT-qPCR was performed with SYBR Green Mix and CFX96 thermocycler (Bio-Rad). The 2^(-ΔΔCt)^ method was used to calculate the relative expression levels (27). Data analysis was performed with the CFX Manager™ Software, Microsoft Excel, and Statistica. RT-qPCR primer sequences are listed in **Table S1**.

### 2.6 Chromatin immunoprecipitation assay

For CHIP assays we used CUT&RUN Assay Kit #86652 (Cell Signaling Technology, Danvers, MA, USA) following the manufacturer’s protocol. Each ChIP assay was repeated 3 times. Briefly, for each reaction we used 100 000 cells. Cells were trypsinized and harvested, washed 3x in ice-cold PBS, bound to concanavalin A-coated magnetic beads for 5 min, RT. Cells were then incubated (4h, 4°C) with 2.5 μg of anti-PHF8 antibody (Abcam, ab36068) or anti-H4K20me1 antibody (Abcam, ab177188) in the antibody-binding buffer plus digitonin that permeabilizes cells. Next, cells are treated with pAG-MNase (1 h, 4°C), washed, and treated with CaCl2 to activate DNA digestion (0.5 h, 4°C). Cells were then treated with the stop buffer and spike-in DNA was added for each reaction for signal normalization, and incubated (10-30 min, 37°C). Released DNA fragments were purified using DNA Purification Buffers and Spin Columns (CS #14209) and quantified by RT-qPCR using primers targeting the promoter, upstream, and downstream regions of the *mTOR* gene (**Table S1**). Rabbit (DA1E) mAb IgG XP® Isotype Control included in the CUT&RUN kit did not afford any signals in RT-qPCR assays targeting *mTOR*.

### 2.7 Confocal microscopy, Aβ staining in N2a-APPswe cells

Mouse neuroblastoma N2a-APPswe cells were cultured in Millicell EZ SLIDE 8-well glass (Merck). After treatments cells were washed 3x with PBS for 10 minutes. Cells were fixed with 4% PFA (Sigma-Aldrich) (37°C, 15 min). After fixation, cells were again washed 3-times with PBS buffer and permeabilized in 0.1% Triton X-100 solution (RT, 20 min), blocked with 0.1% BSA (RT, 1h), and incubated with anti-Aβ antibody (CS #8243; 4°C, 16 h). Cells were then washed 3-times with PBS and stained with secondary antibody Goat Anti-Rabbit IgG H&L (Alexa Fluor® 488) (Abcam, ab150077; RT, 1 h) to detect Aβ. DAPI (Vector Laboratories) was used to visualize nuclei. Fluorescence signals were detected by using a Zeiss LSM 880 confocal microscope with a 488 nm filter for the Alexa Fluor® 488 (Aβ) and 420–480 nm filter for DAPI, taking a *z* stack of 20-30 sections with an interval of 0.66 μm and a range of 15 μm. Zeiss Plan-Apochromat X40/1.2 Oil differential interference contrast objective were used for imaging. Images were quantified with the ImageJ Fiji software (NIH).

### 2.8 Statistical analysis

The results were calculated as mean ± standard deviation. A two-sided unpaired t test was used for comparisons between two groups of variables, *P* < 0.05 was considered significant. Statistical analysis was performed using Statistica, Version 13 (TIBCO Software Inc., Palo Alto, CA, USA, http://statistica.io).

## 3 RESULTS

### 3.1 Cbs deficiency downregulates the histone demethylase Phf8 expression and upregulates H4K20me1 epigenetic mark in the mouse brain

To determine if neuropathy in *Cbs*^-/-^ mice can be caused by the interaction between *Cbs* and *Phf8* genes, we quantified Phf8 protein and mRNA in brains of 9-week-old and 1-year-old *Tg-I278T Cbs*^-/-^ mice and *Tg*-*I278T Cbs*^+/-^ sibling controls by using Western blotting and RT-qPCR assays, respectively. We found that the Phf8 protein levels were significantly reduced in brains of *Tg*-*I278T Cbs*^-/-^ mice *vs. Tg*-*I278T Cbs*^+/-^ sibling controls (9-week-old mice: 0.38±0.27 *vs*. 0.89±0.21, *P* = 0.001; 1-year-old mice: 0.47±0.14 *vs*. 0.93±0.23, *P* = 1.E-05; **Figure 1A**).

**Figure 1.**
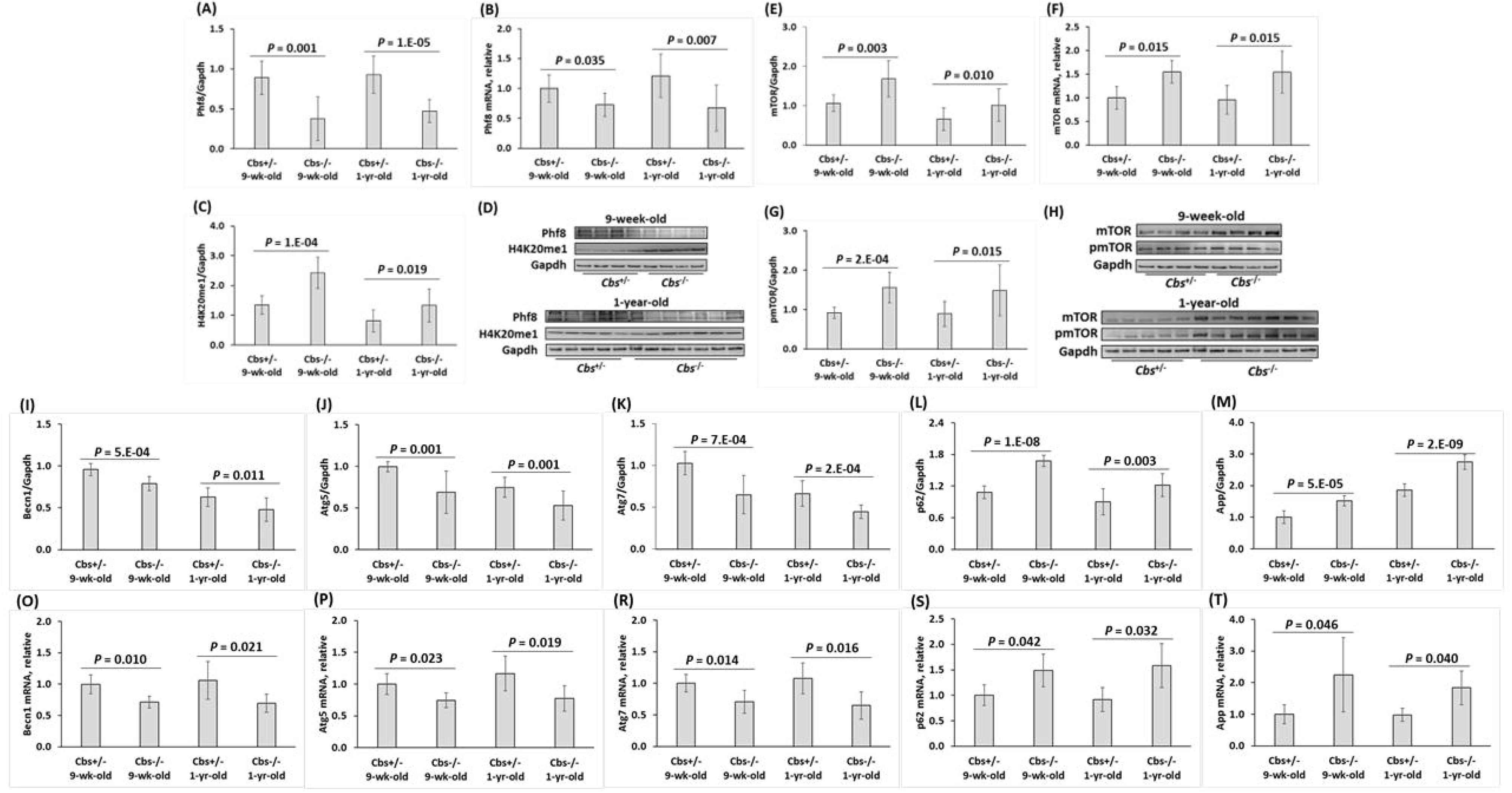
Cbs depletion affects the expression of histone demethylase Phf8, histone H4K20me1 epigenetic mark, mTOR signaling, autophagy, and App in the mouse brain. Nine-week-old and one-year-old *Tg-I278T Cbs*^-/-^ mice (n = 7 and 14) and their *Tg-I278T Cbs*^+/-^ sibling controls (n = 10 and 10) were used in experiments. Bar graphs illustrating quantification of the following brain proteins by Western blotting are shown: Phf8 (**A**), H4K20me1 (**C**), mTOR (**E**), pmTOR (**G**), Becn1 (**I**), Atg5 (**J**), Atg7 (**K**), and p62 (**L**), and App (**M**). Pictures of Western blots used for protein quantification are shown in panels (**D**), (**H**), and (**N**). Bar graphs illustrating quantification of the following brain mRNAs by RT-qPCR are also shown: Phf8 (**B**), mTOR (**F**), Becn1 (**O**), Atg5 (**P**), Atg7 (**R**), and p62 (**S**), and App (**T**). Gapdh protein and mRNA were used as references for normalization.

Phf8 mRNA levels were also significantly reduced in brains of *Tg*-*I278T Cbs*^-/-^ *vs. Tg*-*I278T Cbs*^+/-^ mice (9-week-old mice: 0.73±0.15 *vs*. 1.00±0.15, *P* = 0.035; 1-year-old mice: 0.67±0.26 *vs*. 1.21±0.27, *P* = 0.007; **Figure 1B**). These findings indicate that *Cbs*^-/-^ genotype transcriptionally attenuated Phf8 expression while age did not affect Phf8 levels in *Cbs*^-/-^ nor *Cbs*^+/-^ mice.

The histone H4K20me1 epigenetic mark was significantly elevated in *Tg*-*I278T Cbs*^-/-^ brains compared with *Tg*-*I278T Cbs*^+/-^ brains (9-week-old mice 2.42±0.52 *vs*. 1.34±0.31, *P* = 1.E-04; 1-year-old mice: 1.33±0.56 *vs*. 0.81±0.37, *P* = 0.019; **Figure 1C**). Although *Cbs*^-/-^ genotype upregulated H4K20me1 in 1-year-old mice, age significantly reduced H4K20me1 levels in 1-year-old *vs*. 9-week-old animals (*Cbs*^-/-^: 1.33±0.56 *vs*. 2.42±0.52, *P* = 0.003) and (*Cbs*^+/-^: 0.81±0.37 *vs*. 1.34±0.31, *P* = 1.E-04). These findings show that although age attenuated H4K20me1 levels both in *Cbs*^-/-^ and *Cbs*^+/-^ mice, the effect of *Cbs*^-/-^ genotype on H4K20me1 was not affected by age.

### 3.2 Transcriptional upregulation of mTOR and inhibition of autophagy in *Cbs*^-/-^ mouse brain

We found that mTOR protein was significantly upregulated in brains of *Tg*-*I278T Cbs*^-/-^ *vs. Tg*-*I278T Cbs*^+/-^ mice (9-week-old mice: 1.68±0.46 *vs*. 1.07±0.21, *P* = 0.002; 1-year-old mice: 1.02±0.41 *vs*. 0.66±0.29, *P* = 0.030; **Figure 1E**). mTOR mRNA was also similarly upregulated in *Tg*-*I278T Cbs*^-/-^ *vs. Tg*-*I278T Cbs*^+/-^ mice (9-week-old mice: 1.55±0.24 *vs*. 1.00±0.25, *P* = 0.028; 1-year-old mice: 1.55±0.44 *vs*. 0.96±0.30, *P* = 0.028; **Figure 1F**). While *Cbs*^-/-^ genotype upregulated mTOR in 1-year-old mice, age significantly attenuated mTOR in 1-year-old *vs*. 9-week-old mice (*Cbs*^-/-^: 1.02±0.41 *vs*. 1.68±0.46, *P* = 0.004; *Cbs*^+/-^: 0.66±0.29 *vs*. 1.07±0.21, *P* = 0.002). These findings show that *Cbs*^-/-^ genotype transcriptionally upregulated mTOR in young and old mice while age attenuated mTOR expression both in *Cbs*^-/-^ and *Cbs*^+/-^ mice.

As mTOR is activated by phosphorylation, we quantified mTOR phosphorylated at Ser2448 (pmTOR). We found that pmTOR was significantly elevated in brains of *Tg*-*I278T Cbs*^-/-^ compared to *Tg*-*I278T Cbs*^+/-^ mice (9-week-old mice: 1.56±0.39 *vs*. 0.92±0.14, *P* = 2.E-04; 1-year-old mice: 1.49±0.65 *vs*. 0.90±0.31, *P* = 0.015; **Figure 1G**). These findings indicate that Cbs deficiency upregulated pmTOR to the same extent as mTOR and that age did not affect pmTOR levels in *Cbs*^-/-^ nor *Cbs*^+/-^ mice (*P* = 0.818 and 0.799, respectively).

Because mTOR is a major regulator of autophagy, we quantified autophagy-related proteins in *Tg*-*I278T Cbs*^-/-^ mice. We found that the regulators of autophagosome assembly Becn1, Atg5, and Atg7 were significantly downregulated in brains of 9-week-old *Tg*-*I278T Cbs*^-/-^ mice *vs. Tg*-*I278T Cbs*^+/-^ sibling controls (Becn1: 0.79±0.09 *vs*. 0.96±0.07, *P* = 5.E-04, **Figure 1I;** Atg5: 0.69±0.25 *vs*. 0.99±0.06, *P* = 0.001, **Figure 1J**; Atg7: 0.65±0.23 *vs*. 1.03±0.14, *P* = 7.E-04, **Figure 1K**). *Cbs*^-/-^ genotype significantly downregulated Becn1 (*P* = 0.011), Atg5 (*P* = 0.001), and Atg7 (*P* = 2.E-04) also in 1-year-old mice (**Figure 1I, 1J, and 1K**, respectively). However, Becn1, Atg5, and Atg7 levels were significantly reduced in 1-year-old *vs*. 9-week-old mice, both in *Cbs*^-/-^ mice (Becn1: 0.63±0.11 *vs*. 0.96±0.07, *P* = 3.E-07; Atg5: 0.53±0.17 *vs*. 0.75±0.12, *P* = 0.084; Atg7: 0.45±0.08 *vs*. 0.65±0.23, *P* = 0.007) and *Cbs*^+/-^ animals (Becn1: 0.48±0.14 *vs*. 0.79±0.09, *P* = 4.E-05; Atg5: 0.75±0.12 *vs*. 0.99±0.06, *P* = 1.E-05; Atg7: 0.67±0.15 *vs*. 1.03±0.14, *P* = 1.E-05). The p62 protein, a receptor for degradation of ubiquitinated substrates, was upregulated in 9-week-old *Tg*-*I278T Cbs*^-/-^ mice *vs. Tg*-*I278T Cbs*^+/-^ sibling controls (1.68±0.11 *vs*. 1.08±0.12, *P* = 3.E-08; **Figure 1L**). *Cbs*^-/-^ genotype upregulated p62 also in 1-year-old mice (1.22±0.22 *vs*. 0.90±0.25, *P* = 0.003). However, p62 levels were reduced in 1-year-old *vs*. 9-week-old mice (*Cbs*^-/-^: 1.22±0.22 *vs*. 1.68±0.11, *P* = 5.E-05; *Cbs*^+/-^: 0.90±0.25 *vs*. 1.08±0.12, *P* = 0.056). These findings suggest that autophagy was impaired in *Cbs*^-/-^ mouse brain.

We found similar *Cbs* genotype-dependent changes in autophagy-related mRNAs. Specifically, Becn1, Atg5, and Atg7 (**Figure 2O, P**, and **R**, respectively) were significantly downregulated, while p62 (**Figure 2S**), were significantly upregulated in *Tg*-*I278T Cbs*^-/-^ compared with *Tg*-*I278T Cbs*^+/-^ mice, reflecting changes in the corresponding protein levels. These findings indicate that Cbs gene exerts transcriptional control over the expression of mTOR, Becn1, Atg5, Atg7, and p62 in the mouse brain.

**Figure 2.**
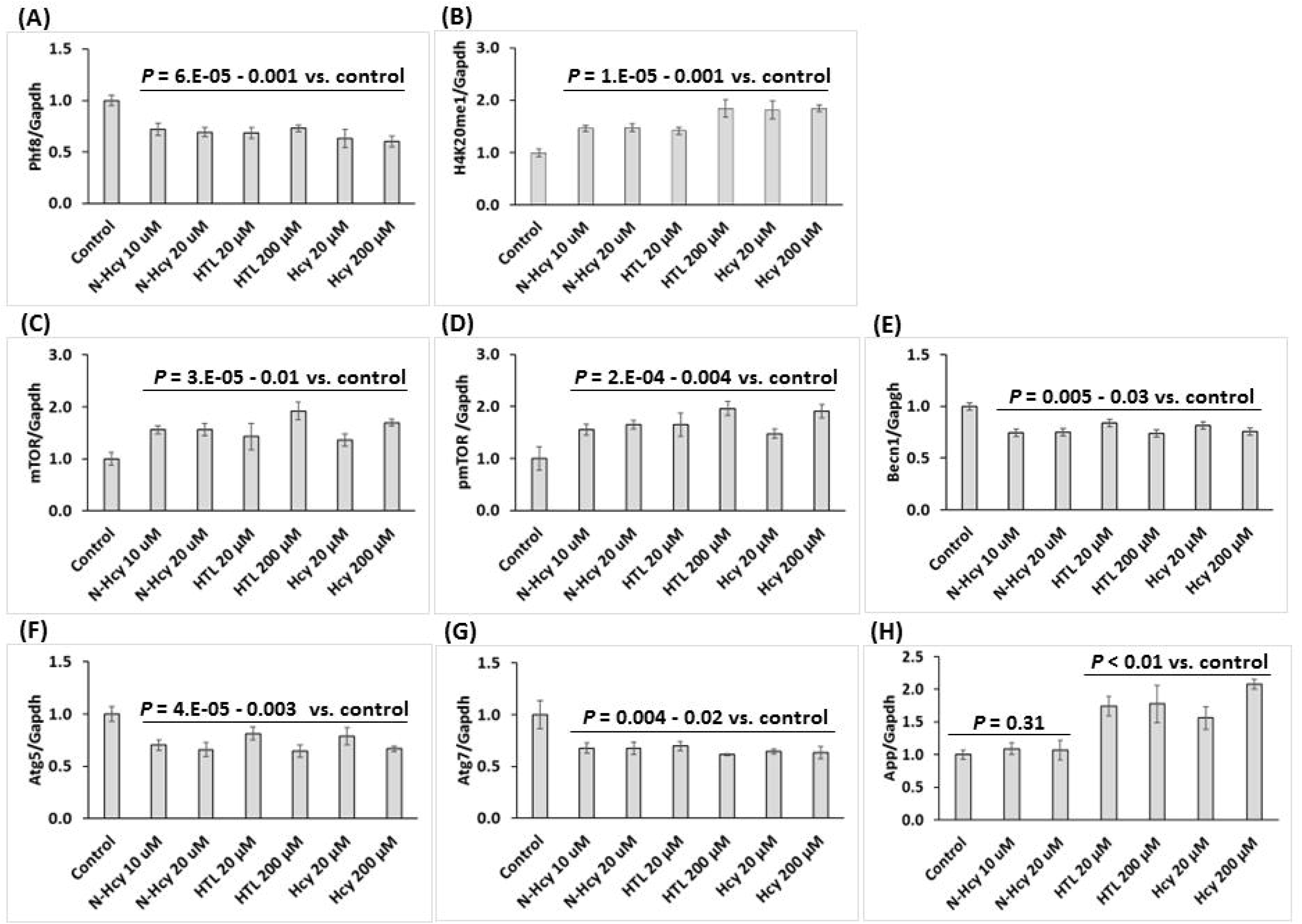
Hcy-thiolactone, *N*-Hcy-protein, and Hcy downregulate Phf8, upregulate H4K20me1 epigenetic mark, mTOR signaling, App, and impair autophagy in mouse neuroblastoma N2a cells. N2a cells were treated with indicated concentrations of *N*-Hcy-protein, Hcy-thiolactone (HTL), or Hcy for 24 h at 37°C as described in Materials and Methods. Bar graphs illustrating the quantification of Phf8 (**A**), H4K20me1 (**B**), mTOR (**C**), pmTOR (**D**), App (**E**), Becn1 (**F**), Atg5 (**G**), and Atg7 (**H**) based on Western blots analyses with corresponding antibodies are shown. Gapdh was used as a reference protein. Data are averages of three independent experiments.

### 3.3 Cbs deficiency upregulates App expression in mouse brain

We have previously found that Aβ was elevated in brains of 1-year-old mice *Cbs*^-/-^ mice compared to *Cbs*^+/-^ controls (28). To determine whether upregulated App expression might be responsible for elevated Aβ, we quantified App protein and mRNA in *Tg*-*I278T Cbs*^-/-^ and *Tg*-*I278T Cbs*^+/-^ mice by Western blotting (**Figure 1M**) and RT-qPCR (**Figure 1T**), respectively.

We found that App protein was significantly elevated in 9-week-old *Tg*-*I278T Cbs*^-/-^ *vs. Tg*-*I278T Cbs*^+/-^ mice (1.52-fold: 1.52±0.16 *vs*. 1.01±0.20, *P*_*genotype*_ = 5.E-05). We found a similar elevation in App protein in 1-year-old *Tg*-*I278T Cbs*^-/-^ *vs. Tg*-*I278T Cbs*^+/-^ mice (1.49-fold, 2.75±0.23 *vs*. 1.85±0.20, *P*_*genotpe*_ = 2.E-09) (**Figure 1M**). App mRNA was also upregulated in *Tg*-*I278T Cbs*^-/-^ *vs. Tg*-*I278T Cbs*^+/-^ mice (9-week-old mice: 2.25±1.17 *vs*. 1.00±0.30, *P* = 0.046; 1-year-old mice: 1.84±0.54 *vs*. 0.98±0.21, *P* = 0.020; **Figure 1T**). These findings indicate that Cbs exerts transcriptional control over App expression.

We also found that App protein significantly increased with age (**Figure 1M**), irrespective of Cbs genotype: old/young *Tg*-*I278T Cbs*^-/-^ mice 1.80-fold, *P*_*age*_ = 1.E-10; old/young *Tg*-*I278T Cbs*^+/-^ mice 1.85-fold, *P* = 3.E-08. However, age did not affect App mRNA levels (**Figure 1T**). These findings suggest that age exerts translational control over App expression independent of *Cbs* genotype or that age exerts posttranslational control by reducing degradation of App protein.

### 3.4 *Cbs* gene silencing downregulates the histone demethylase Phf8, upregulates H4K20me1, mTOR, pmTOR, APP, and inhibits autophagy in mouse neuroblastoma N2A-APPswe cells

To elucidate the mechanism by which Cbs deficiency impacts Phf8 and its downstream effects on mTOR, autophagy, and App we first examined whether the findings in *Cbs*^-/-^ mice can be recapitulated in cultured mouse neuroblastoma N2A-APPswe cells. We silenced the *Cbs* gene by transfecting corresponding siRNAs into N2A-APPswe cells and studied how the silencing impacts the level of Phf8 and its downstream effects. Changes in individual mRNA and protein levels in *Cbs*-silenced cells were analyzed by RT-qPCR and Western blotting, respectively, using Gapdh mRNA and protein as references.

We found that the Cbs protein level was reduced by 80% in *Cbs*-silenced cells (*P* = 2.E-09; **Figure S1A**). We also found that the histone demethylase Phf8 protein level was also significantly reduced (by 39%, *P* = 2.E-06; **Figure S1B**), while the histone H4K20me1 level was significantly elevated (2-fold, *P* = 4.E-05; **Figure S1C**) in *Cbs*-silenced N2a-APPswe cells.

At the same time, mTOR protein was significantly upregulated in *Cbs*-silenced N2a-APPswe cells (1.7-fold, *P* = 1.E-05; **Figure S1D)**, as was pmTOR (2-fold, *P* = 3.E-08; **Figure S1E**) and APP (1.5-fold, *P* = 7.E-07; **Figure S1F**), while autophagy-related proteins Becn1, Atg5, and Atg7 (**Figure S1G, S1H**, and **S1I**, respectively) were significantly downregulated (by 30-40%, *P* = 4.E-08 to 1.E-06).

We found similar changes in levels of mRNAs in *Cbs*-silenced N2a-APPswe cells (**Figure S2**). Specifically, Cbs mRNA level was reduced (by 85%, *P* = 2.E-04; **Figure S2A**), as were Phf8 mRNA (by 50%, *P* = 0.001; **Figure S2B**) and mRNAs for autophagy-related proteins Becn1, Atg5, and Atg7 (by 30-40%, *P* = 0.002 – 0.023; **Figure S2D, S2E**, and **S2F**, respectively). mTOR mRNA was significantly upregulated (1.5-1.7-fold, *P* = 0.019; **Figure S2C**) as was APP mRNA (1.7-1.9-fold, *P* = 0.005 – 0.021; **Figure S2G**) in *Cbs*-silenced N2a-APPswe cells, reflecting changes in the corresponding protein levels (**Figure 1A-N**). These findings indicate that *Cbs* gene exerts transcriptional control over the expression of Phf8, mTOR, App, Becn1, Atg5, and Atg7.

The Western blot and RT-qPCR results show that the changes in Phf8, H4K20m31, mTOR signaling, autophagy, and APP induced in N2a-APPswe cells by *Cbs* gene silencing (**Figure S1** and **S2**) recapitulate the *in vivo* findings in the *Cbs*^-/-^ mouse brain (**Figure 1**).

### 3.5 Hcy-thiolactone, *N*-Hcy-protein, and Hcy downregulate the histone demethylase Phf8, upregulate the H4K20me1 epigenetic mark and mTOR, and impair autophagy in N2a cells

Because Hcy-thiolactone and *N*-Hcy-protein (14-17), in addition to Hcy (29), are elevated in *Cbs*^-/-^ mice, it is difficult to assign observed phenotypes to a specific metabolite in these mice. However, this limitation can be overcome in cultured cells by treatments with an excess of a specific metabolite. To determine how each metabolite affects the expression of Phf8 and its effects on downstream targets, we treated N2a cells with Hcy-thiolactone, *N*-Hcy-protein, and Hcy.

We found significantly reduced Phf8 levels in N2a cells treated with 20 μM Hcy-thiolactone (0.69±0.05), Hcy (0.63±0.09), or 10 μM *N*-Hcy-protein (0.72±0.06) compared to control (1.00±0.05, *P* < 0.001; **Figure 2A**).

In contrast, levels of H4K20me1 mark were significantly elevated in N2a cells treated with 20 μM Hcy-thiolactone (1.42±0.07), Hcy (1.81±0.17), or 10 μM *N*-Hcy-protein (1.46±0.05) compared to untreated control cells (1.00±0.07, *P* < 0.001; **Figure 2B**).

mTOR levels were significantly upregulated in N2a cells by treatments with these metabolites: 20 μM Hcy-thiolactone (1.46±0.25), Hcy (1.36±0.12), or 10 μM *N*-Hcy-protein (1.56±0.08) compared to untreated control cells (1.00±0.12, *P* < 0.001; **Figure 2C**). Levels of pmTOR were also significantly upregulated by these treatments: 20 μM Hcy-thiolactone (1.65±0.22-fod), Hcy (1.48±0.08-fold), 10 μM *N*-Hcy-protein (1.56±0.11-fold) compared to control (1.00±0.12, *P* < 0.001; **Figure 2D**).

App levels were also significantly upregulated in N2a cells treated with 20 μM Hcy-thiolactone (1.74-fold, *P* = 0.002) or Hcy (1.56-fold, *P* = 0.002 (**Figure 2E**). Increasing Hcy to 200 μM significantly increased upregulation of App from 1.56- to 2.08-fold (*P* = 0.002) while higher concentration of Hcy-thiolactone did not further elevate App levels (**Figure 2E**).

Autophagy-related proteins Becn1, Atg5, and Atg7 were significantly downregulated (by 16-25%, *P* < 0.001) in N2a cells treated with Hcy-thiolactone, *N*-Hcy-protein, or Hcy (**Figure 2F, 2G, and 2H**, respectively).

Similar effects were observed in N2a cells treated with 10-fold higher concentrations of Hcy-thiolactone and Hcy (200 μM), or 2-fold higher *N*-Hcy-protein (20 μM) (**Figure 2**). These findings show that Hcy and its downstream metabolites Hcy-thiolactone and *N*-Hcy-protein, which are elevated in *Cbs*^-/-^ mice (14-17), each can affect the expression of Phf8 and its downstream targets.

### 3.6 *Cbs* gene silencing increased H4K20me1 biding to mTOR promoter in N2a cells

To determine whether increased levels of the histone H4K20me1 mark can promote mTOR gene expression by binding to its promoter, we carried out ChIP experiments using anti-H4K20me1 antibody (**Figure 3**). The *Cbs* gene was silenced by transfecting N2a using two different siRNAs. The cells were permeabilized, treated with anti-H4K20me1 antibody and a recombinant micrococcal nuclease-protein A/G. DNA fragments released form N2a-APPswe cells we quantified by RT-qPCR using primers targeting the transcription start site (TSS) of the mTOR gene as well as upstream (UP) and downstream (DOWN) regions from the TSS. We found that in siRNA *Cbs*-silenced N2a-APPswe cells the binding of H4K20me1 was significantly increased at the mTOR TSS (2.7-fold, *P* = 0.002), UP (1.7-fold, *P* = 0.013), and DOWN (2.2-fold, *P* = 3.E-04) sites (**Figure 3A**). Importantly, there were significantly more DNA fragments from the TSS site (2.62±0.31 and 2.68±0.19 for siRNA Cbs #1 and #2, respectively) than from the UP (1.67±0.14 to 1.68±0.41 for siRNA Cbs #1 and #2, respectively; *P* = 1.E-04) and DOWN (2.22±0.06 to 2.27±0.19 for siRNA Cbs #1 and #2, respectively; *P* = 5.E-04) sites (**Figure 3A**). Control experiments show that the binding of H3K4me3 to RPL30 intron was not affected by *Cbs* gene silencing (**Figure 3B**). These findings indicate that the binding of H4K20me1 was significantly higher at TSS than at UP and DOWN sites in siRNA *Cbs*-silenced cells. Similar results were obtained with N2a-APPswe cells (not shown).

**Figure 3.**
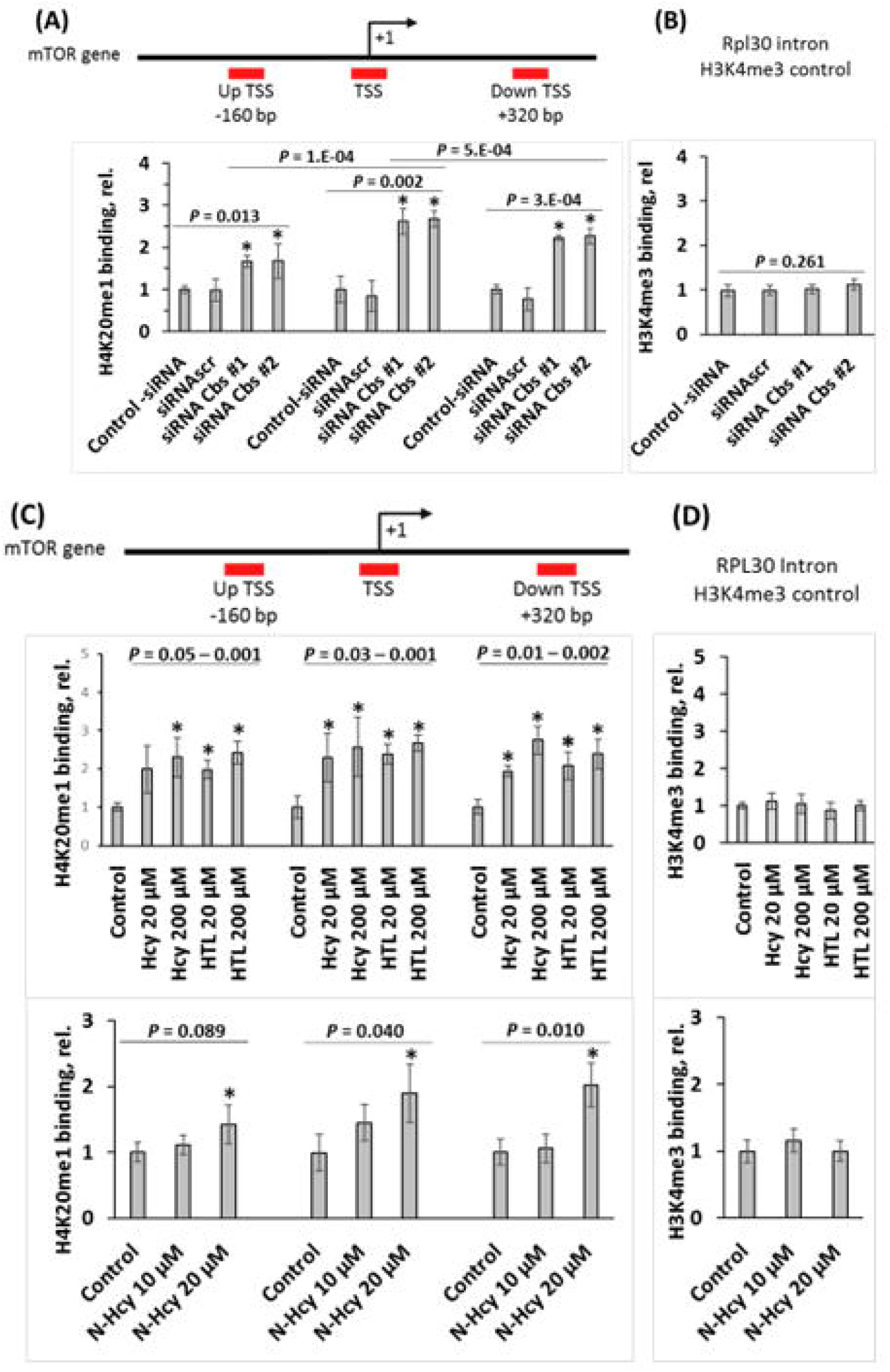
*Cbs* gene silencing or treatments with Hcy-thiolactone, *N*-Hcy-protein, and Hcy increase H4K20me1 binding at the *mTOR* promoter in mouse neuroblastoma N2a cells. (**A**) CHIP assays with anti-H4K20me1 antibody show the specific binding of H4K20me1 at the transcription start site (TSS) of the *mTOR* gene as well as downstream and upstream sites in Cbs siRNA silenced N2a cells. Bar graphs show the relative H4K20me1 binding at indicated regions of the *mTOR* gene in N2a cells transfected with two different siRNAs targeting the *Cbs* gene (siRNA Cbs #1 and #2). Transfections without siRNA (Control -siRNA) or with scrambled siRNA (siRNAscr) were used as controls. (**B**) Control CHIP experiment with anti-H3K4me3 antibody shows that *Cbs* gene-silencing did not affect the binding of H3K4me3 at the Rpl30 intron. RT-qPCR was carried out on the input and precipitated DNA fragments. (**C**) N2a cells were treated with indicated concentrations of *N*-Hcy-protein, Hcy-thiolactone (HTL), or Hcy for 24h at 37°C. Untreated cells were used as controls. CHIP assays with anti-H4K20me1 antibody show the binding of H4K20me1 at the transcription start site (TSS) of the *mTOR* gene as well as downstream and upstream sites. Bar graphs show the relative H4K20me1 binding at the indicated regions of the *mTOR* gene. (**D**) Control CHIP assay with anti-H3K4me3 antibody shows that Hcy-thiolactone, *N*-Hcy-protein, and Hcy did not affect the binding of H3K4me3 at the Rpl30 intron. RT-qPCR was carried out on the input and precipitated DNA fragments. * Significant difference *vs*. control, *P* < 0.05.

CHIP experiments with anti-Phf8 antibody showed that Cbs silencing did not affect binding of Phf8 to the mTOR gene (not shown).

### 3.7 Hcy-thiolactone, *N*-Hcy-protein, and Hcy increase H4K20me1 binding to mTOR promoter in N2a cells

Hcy-thiolactone and *N*-Hcy-protein (14-17) are elevated in *Cbs*^-/-^ mice, as is Hcy (29). Each of these metabolites can affect mTOR expression by promoting H4K20me1 binding at the mTOR promoter. To examine this possibility, we treated N2a-APPswe cells with Hcy-thiolactone, *N*-Hcy-protein, or Hcy for 24 h and analyzed how these treatments affect levels of H4K20me1 bound to the mTOR gene using ChIP experiments with anti-H4K20me1 antibody. We found that Hcy-thiolactone at 20 μM increased binding of H4K20me1 at the mTOR TSS (2.4-fold, *P* = 0.004), UP (2.0-fold, *P* = 0.003), and DOWN (2.1-fold, *P* = 0.011) sites (**Figure 3C**). Similar results were obtained with 200 μM Hcy-thiolactone.

Hcy at 20 μM increased binding of H4K20me1 at the mTOR TSS (2.3-fold, *P* = 0.032), UP sites (2.3-fold, *P* = 0.052), and DOWN sites (1.9-fold, *P* = 0.002). Hcy at 200 μM significantly increased binding of H4K20me1 at the mTOR TSS (2.6-fold, *P* = 0.030), UP (2.4-fold, *P* = 0.013), and DOWN sites (2.8-fold, *P* = 0.002) TSS (**Figure 3C**). Similar results were obtained with N2a-APPswe cells (not shown).

*N*-Hcy-protein at 20 μM significantly increased binding of H4K20me1 at the *mTOR* TSS (1.9-fold, *P* = 0.040) and DOWN (2.0-fold, *P* = 0.010) sites, but not UP site (1.4-fold, *P* = 0.089) (**Figure 3C**). Smaller, nonsignificant increases were observed with 10 μM *N*-Hcy-protein.

Neither Hcy, Hcy-thiolactone, nor *N*-Hcy-protein affected binding of Phf8 at the *mTOR* TSS, UP and down sites of the mTOR gene (not shown).

Control experiments show that H3K4me3 binding to RPL30 intron was not affected by Hcy, Hcy-thiolactone nor *N*-Hcy-protein (**Figure 3D**). These findings indicate that Hcy-thiolactone significantly increased binding of H4K20me1 at the mTOR gene TSS, UP, and DOWN sites while Hcy significantly increased binding of H4K20me1 at the mTOR TSS.

### 3.8 *Cbs* gene silencing upregulates Aβ in N2a-APPswe cells

To determine whether Cbs depletion could affect Aβ accumulation, we silenced the *Cbs* gene in N2a-APPswe cells and quantified Aβ by fluorescence confocal microscopy using anti-Aβ antibody. The cells were transfected with two different *Cbs*-targeting siRNAs, permeabilized, treated with anti-Aβ antibody, and Aβ was visualized with fluorescent secondary antibody (**Figure 4A**) and quantified (**Figure 4B**). We found that *Cbs*-silencing led to increased Aβ accumulation, manifested by significant increases in the area (from 130±7 to 189±47 μm^2^ and 183±13 μm^2^ for siRNA Cbs #1 and #2, respectively; *P* = 0.007) and an average size of fluorescent Aβ puncta (from 0.32±0.02 to 0.80±0.02 μm^2^ and to 0.90±0.06 μm^2^ for siRNA Cbs #1 and #2, respectively; *P* = 1.E-06) in *Cbs* siRNA-treated N2a-APPswe cells compared to siRNAscr-treated cells or control without siRNA (**Figure 4B**).

**Figure 4.**
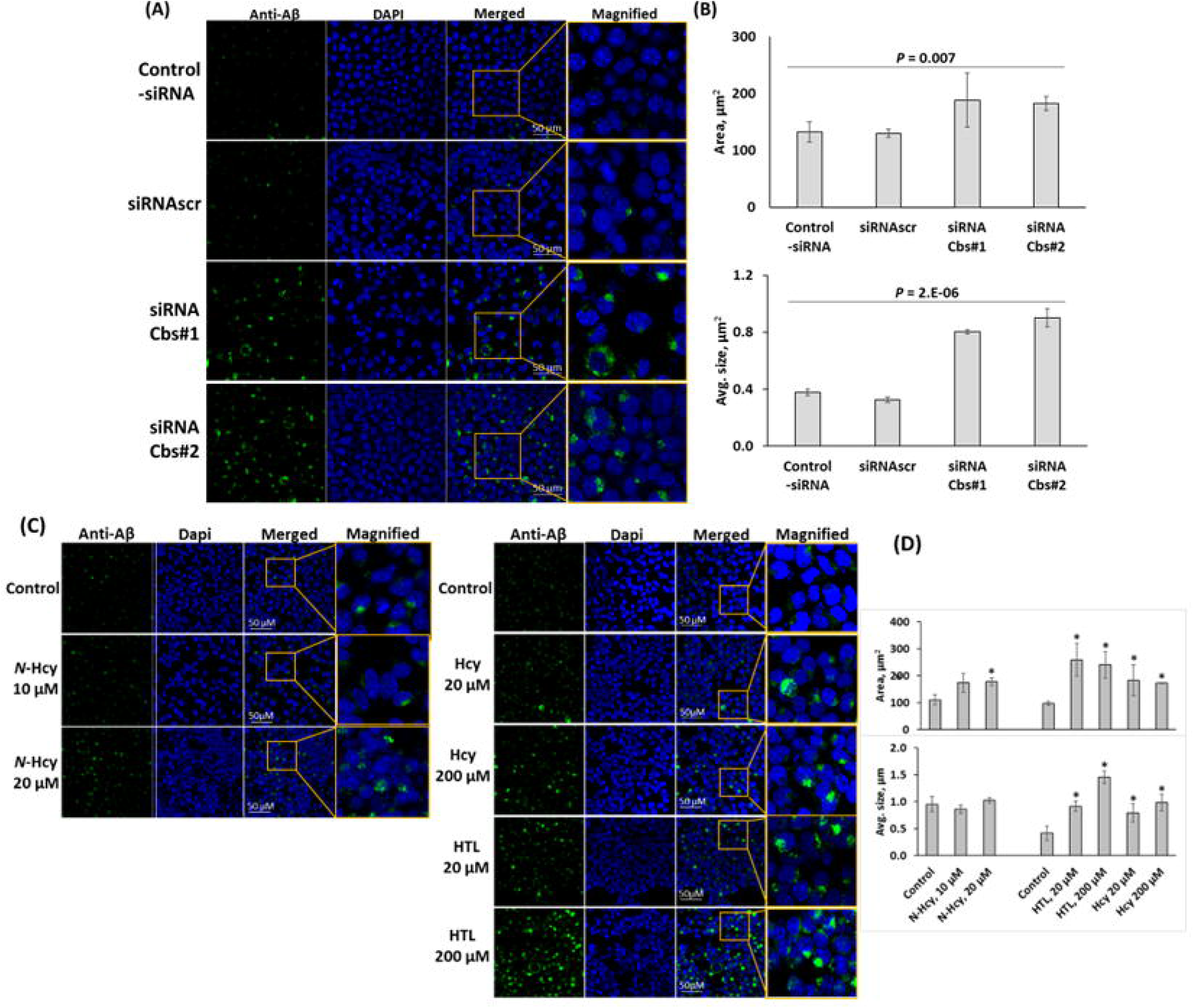
Cbs depletion or treatments with Hcy-thiolactone, *N*-Hcy-protein, and Hcy promote Aβ accumulation in the mouse neuroblastoma N2a-APPswe cells. (**A, B**) The cells were transfected with siRNAs targeting the *Cbs* gene. Transfections without siRNA (Control -siRNA) or with scrambled siRNA (siRNAscr) were used as controls. Aβ was detected and quantified by confocal immunofluorescence microscopy using anti-Aβ antibody. (**A**) Confocal microscopy images of Aβ signals from *Cbs*-silenced N2a-APPswe cells. (**B**) Bar graphs show quantification of Aβ signals in *Cbs*-silenced and control cells. (**C, D)** *N*-Hcy-protein, Hcy-thiolactone (HTL), or Hcy promote Aβ accumulation in mouse neuroblastoma N2a-APPswe cells. N2a-APPSwe cells were treated with indicated concentrations of *N*-Hcy-protein, Hcy-thiolactone (HTL), or Hcy for 24h at 37°C. Untreated cells were used as controls. Confocal microscopy images (**C**) and quantification of Aβ signals (**D**) from treated and untreated N2a-APPswe cells. Data for cells treated with Hcy-thiolactone and *N*-Hcy-protein in panels (**C, D)** were reproduced with permission from ref. (30).

### 3.9 Hcy-thiolactone, *N*-Hcy-protein, and Hcy upregulate Aβ accumulation in N2a-APPswe cells

To ascertain how each Hcy metabolite can affect Aβ accumulation, we treated N2a-APPswe cells with Hcy-thiolactone, *N*-Hcy-protein (30), or Hcy and quantified Aβ by fluorescence confocal microscopy using anti-Aβ antibody (**Figure 4C, D**). The metabolite-treated and untreated control cells were permeabilized, treated with anti-Aβ antibody, followed by fluorescent secondary antibody to visualize Aβ signals. We found that Aβ was significantly upregulated in N2a-APPswe cells treated with 20 μM Hcy, Hcy-thiolactone, or 10 μM *N*-Hcy-protein, manifested by significantly increased average size and signal intensity of the fluorescent Aβ puncta, compared to untreated cells (**Figure 4D**). Similar results were obtained with N2a-APPswe cells treated with 10-fold higher concentrations of Hcy, Hcy-thiolactone (200 μM), or 2-fold higher *N*-Hcy-protein (20 μM) (30). However, while treatments with Hcy or Hcy-thiolactone increased the size of the fluorescent Aβ puncta, treatments with *N*-Hcy-protein did not affect the size of Aβ signal (**Figure 4D**), suggesting different effects of these metabolites on the structure of Aβ aggregates. These findings indicate that Hcy and its downstream metabolites Hcy-thiolactone and *N*-Hcy-protein contribute to increased accumulation of Aβ induced by the metabolic stress of HHcy.

### 3.9 *Phf8* gene silencing upregulates Aβ but not APP in N2a-APPswe cells

The findings that Phf8 expression was attenuated in brains of *Cbs*^-/-^ mice and in *Cbs*-silenced or Hcy metabolite-treated mouse neuroblastoma N2a-APPswe cells raises a possibility that Phf8 loss by itself can affect biochemical pathways leading to Aβ accumulation. To examine this possibility, we silenced the *Phf8* gene by transfection of N2a-APPswe cells with Phf8-targeting siRNAs (31) and quantified by Western blotting proteins that were affected by Cbs depletion in the mouse brain (**Figure 1**). We found significantly reduced Phf8 levels (by 80%, *P* = 7.E-09; **Figure 5A**), significantly increased H4K20me1 (3-fold, *P* = 7.E-09; **Figure 5B**), mTOR (1.4-fold, *P* = 7.E-06; **Figure 5C**), and pmTOR (1.6-fold, *P* = 5.E-04; **Figure 5D**) levels in Phf8-silenced cells. Autophagy-related proteins Atg5 and Atg7 were significantly downregulated (by 20-35%, *P* = 3.E-04 – 2.E-06; **Figure 5E, F**) while Becn1 was not affected in Phf8-silenced cells (**Figure 5G**).

**Figure 5.**
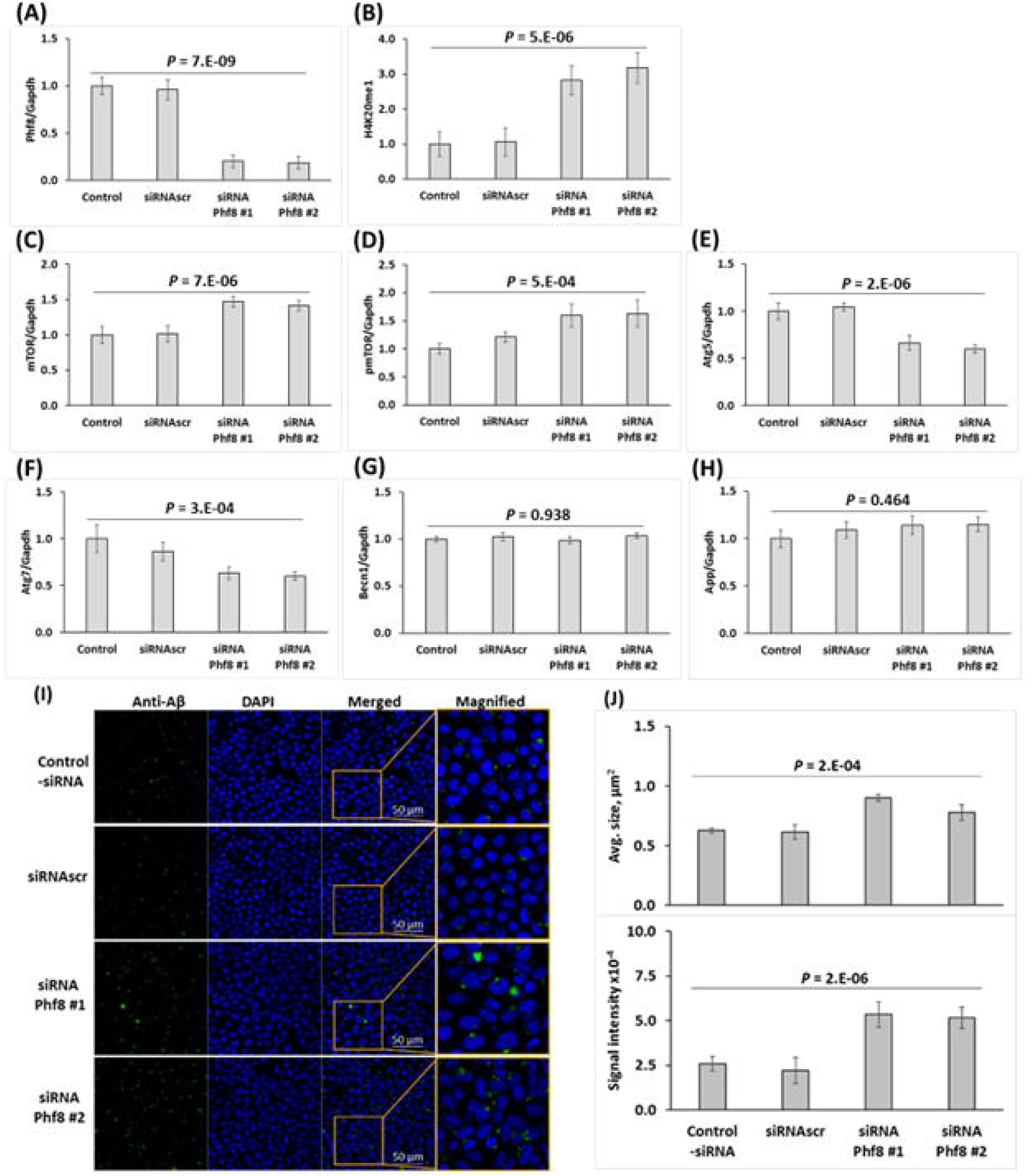
Silencing *Phf8* gene promotes Aβ accumulation mediated by upregulation of mTOR signaling and inhibition of autophagy in the mouse neuroblastoma N2a-APPswe cells. The cells were transfected with siRNAs targeting the *Phf8* gene (Phf8 siRNA #1 and #2). Transfections without siRNA (Control - siRNA) or with scrambled siRNA (siRNAscr) were used as controls. Proteins were quantified by Western blotting. Bar graphs illustrate levels of **(A)** Phf8, **(B)** H4K20me1, **(C)** mTOR, **(D)** pmTOR, **(E)** Atg5, **(F)** Atg7, **(G)** Becn1, and **(H)** App. Aβ was detected and quantified by confocal immunofluorescence microscopy using anti-Aβ antibody. (**I**) Confocal microscopy images of Aβ signals from *Cbs*-silenced and control N2a-APPswe cells. (**J**) Bar graphs show quantification of Aβ signals. Reproduced with permission from ref. (31).

Importantly, Western blot analyses showed that the *Phf8* gene silencing did not affect APP levels in N2a-APPswe cells (**Figure 5H**). In contrast, fluorescence confocal microscopy analyses showed that Aβ was significantly upregulated in *Phf8*-silenced cells, manifested by significantly increased average size (*P* = 2.E-04) and signal intensity (*P* = 2.E-06) of the fluorescent Aβ puncta compared to controls without siRNA or with siRNAscr (**Figure 5I, J**) (31). Taken together, these findings clearly show that Aβ accumulation in the *Phf8*-silenced cells occurred independently of APP and suggest that most likely it was caused by impaired autophagy.

## 3 DISCUSSION

The loss of CBS, an important Hcy-metabolizing enzyme (12, 13), or the loss of PHF8 (21, 22), an important histone demethylase participating in epigenetic regulation, causes severe mental retardation in humans. Similar neuropathies are also observed in *Cbs*^-/-^ and *Phf8*^-/-^ mice. Because PHF8 is involved in epigenetic regulation, we surmised that dysregulation of epigenetic mechanisms involving PHF8 could underlie neuropathy associated with CBS deficiency. The present study substantiated this supposition by showing that the expression of Phf8 mRNA and protein was downregulated while H4K20me1 epigenetic mark was upregulated in brains of *Cbs*^-/-^ mice and in Cbs-depleted mouse neuroblastoma cells (**Figure 6**).

**Figure 6.**
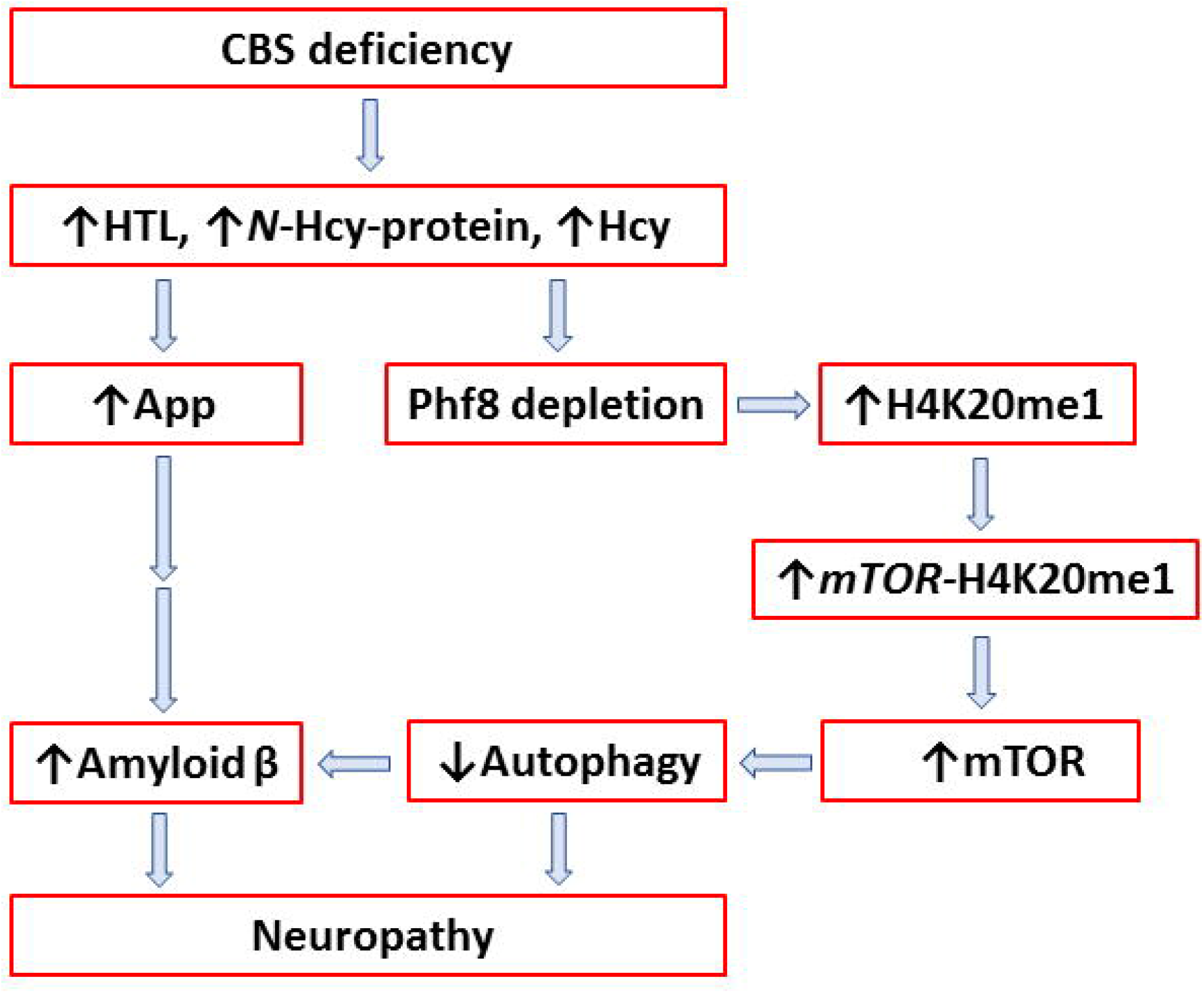
Mechanisms underlying neuropathy in the Cbs-deficient brain. Up and down arrows show direction of changes in the indicated metabolites, proteins, and molecular processes. CBS, cystathionine β-synthase; Hcy, homocysteine; HTL, Hcy-thiolactone; App, amyloid beta precursor protein; mTOR, mammalian target of rapamycin; Phf8, Plant Homeodomain Finger protein 8.

We also found that metabolites that are elevated in CBS-deficient humans and mice – Hcy-thiolactone, *N*-Hcy-protein, and Hcy – downregulated Phf8 (**Figure 2A**) and upregulated histone H4K20me1 (**Figure 2B**) in cultured mouse neuroblastoma cells. This suggests that Hcy-thiolactone, *N*-Hcy-protein, and Hcy are responsible for Phf8 downregulation (**Figure 1A**) and H4K20me1 upregulation (**Figure 1B**) found *in vivo* in brains of *Cbs*^-/-^ mice. We also showed that treatments with siRNA targeting *Cbs* gene or with Hcy-thiolactone or Hcy, which metabolically downregulated Phf8 expression (**Figure S1B, Figure 2A**), promoted the accumulation of both APP (**Figure 2H**) and Aβ (**Figure 4**) in mouse neuroblastoma cells. In contrast, attenuation of Phf8 expression by treatments with siRNAs targeting *Phf8* gene, promoted Aβ accumulation (**Figure 5I, J**), but did not affect App levels (**Figure 5H**). These disparate effects of metabolic and genetic Phf8 depletion on App expression suggest that Aβ accumulation can occur by two different mechanisms in Cbs-depleted brain and neural cells: upregulated App expression caused by Hcy-thiolactone, *N*-Hcy-protein, and Hcy, and impaired Aβ clearance caused by downregulation of autophagy (**Figure 6**). In Phf8-depleted cells, only one mechanism, downregulated autophagy, leads to accumulation of Aβ.

Our findings suggest that Phf8 regulates Aβ accumulation through its effects on mTOR and autophagy. Specifically, we found that treatments with Hcy-thiolactone, *N*-Hcy-protein, or Hcy, which downregulated Phf8 expression (**Figure 2A**) and upregulated the histone mark H4K20me1 (**Figure 2B**), also increased H4K20me1 binding to the mTOR promoter in mouse neuroblastoma cells (**Figure 3C**). These findings provide direct mechanistic evidence linking each of the Hcy metabolites with dysregulated mTOR signaling and its downstream consequences. Concomitant inhibition of autophagy (**Figure 2 F, G, H**) and upregulation of Aβ (**Figure 4C, D**) by Hcy-thiolactone, *N*-Hcy-protein, and Hcy identify a likely mechanism that can contribute to the neuropathy of CBS deficiency (**Figure 6**) and can also account for the association of HHcy with Alzheimer’s disease (18).

Previous studies showed that Phf8 bound at the TSS of *Kras, Camk2d*, and *Rps6ka1* genes involved in mTOR signaling and that depletion of Phf8 increased binding of H4K20me1 at the TSS in *Kras, Camk2d*, and *Rps6ka1* genes in mice (22); H4K20me1 binding at the *mTOR* gene was not examined in that study. In the present study we found that Phf8 depletion in mouse neuroblastoma N2a cells by treatments with Hcy-thiolactone, *N*-Hcy-protein, or Hcy (**Figure 2A**), or by silencing *Cbs* gene with siRNAs (**Figure S2B**), increased H4K20me1 binding at the TSS as well as down and up sites of the mTOR gene (**Figure 3A, 3C**). Our findings add mTOR to the list of genes regulated by H4K20me1.

However, in contrast to TSS of the *Kras, Camk2d*, and *Rps6ka1* genes, which bound Phf8 (22), we could not detect any binding of Phf8 to TSS of the *mTOR* gene.

Previous studies have found that *Cbs*^-/-^ mice have elevated Hcy-thiolactone and *N*-Hcy-protein (14-17), in addition to Hcy (29), which was accompanied by increased brain Aβ accumulation (28) and cognitive impairments (12, 13). In the present study we have found significantly depleted Phf8 in brains of *Cbs*^-/-^ mice (**Figure 1A**). Although depletion of Phf8 is linked to intellectual disability, autism spectrum disorder, attention deficit hyperactivity disorder (20), and mental retardation (21), it was not known to be associated with Alzheimer’s disease. Our present findings that Phf8 depletion in the mouse neuroblastoma cells, induced by *Phf8* siRNA interference (**Figure 5A**), or by supplementation with Hcy-thiolactone, *N*-Hcy-protein, or Hcy (**Figure 2A**), significantly increased Aβ accumulation (**Figure 5I, J**), suggest that Phf8 depletion can contribute to the association of Alzheimer’s disease with HHcy (18).

In conclusion, our findings identify the histone demethylase Phf8 as a regulator of Aβ accumulation and suggest that neuropathy of CBS deficiency is mediated by Hcy metabolites-dependent depletion of Phf8, which upregulates mTOR through increased H4K20me1 binding at the mTOR promoter, which in turn inhibits autophagy, and leads to Aβ accumulation. We have also shown that Cbs-depletion and Hcy metabolites, independently of their inhibitory effect on Phf8 expression, upregulated App expression, which can also contribute to Aβ accumulation.

## AUTHOR CONTRIBUTIONS

Ł. Witucki performed and analyzed the experiments; H. Jakubowski conceived the idea for the project, designed the study, bred the mice, collected mouse tissue samples, analyzed data, and drafted the paper.

## ACKNOWLEDGMENTS

We thank S. S. Sisodia for kindly providing mouse neuroblastoma cells expressing human APP-695 harboring the K670N/M671L Swedish double mutation associated with familial early-onset Alzheimer’s disease, N2a-APPswe. Supported in part by grants 2018/29/B/NZ4/00771, 2019/33/B/NZ4/01760, and 2021/43/B/NZ4/00339 from the National Science Center, Poland.

## DATA AVAILABILITY STATEMENT

The data that support the findings of this study are available in the methods and/or supplementary material of this article.

