## Supplementary for "Homocysteine Metabolites Inhibit Autophagy and Elevate Amyloid Beta by Impairing Phf8/H4K20me1-dependent Epigenetic Regulation of mTOR in Cystathionine β-Synthase-Deficient Mice"

**Supplementary Material**

Supplementary **Figure S1**

Supplementary **Figure S2**

Supplementary **Table S1**

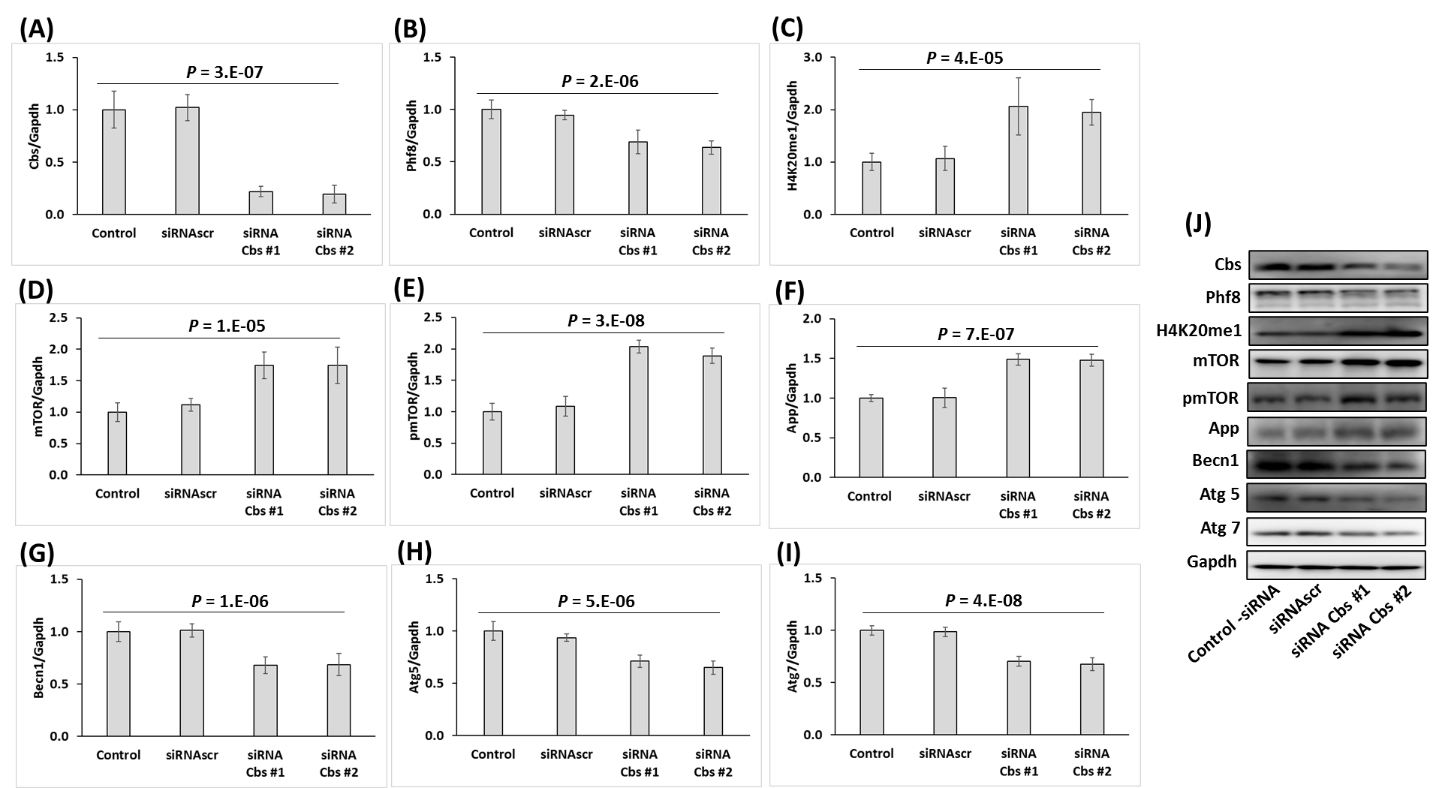

**Figure S1**. *Cbs* gene silencing in mouse neuroblastoma N2a-APPswe cells recapitulates changes in histone demethylase Phf8, H4K20me1, mTOR signaling, App, and autophagy-related protein levels observed in *Cbs*^-/-^ mouse brain. Bar graphs illustrating the quantification of Cbs (**A**), Phf8 (**B**), H4K20me1 (**C**), mTOR (**D**), pmTOR (**E**), Bcln1 (**F**), Atg5 (**G**), Atg7 (**H**), and App (**I**) in N2a-APPswe cells transfected with two different siRNAs targeting the *Cbs* gene (siRNA Cbs #1 and #2) are shown. Transfections without siRNA (Control -siRNA) or with scrambled siRNA (siRNAscr) were used as controls. Representative Western blots are shown in panel (**J**). Gapdh was used as a reference protein. Data are averages of three independent experiments.

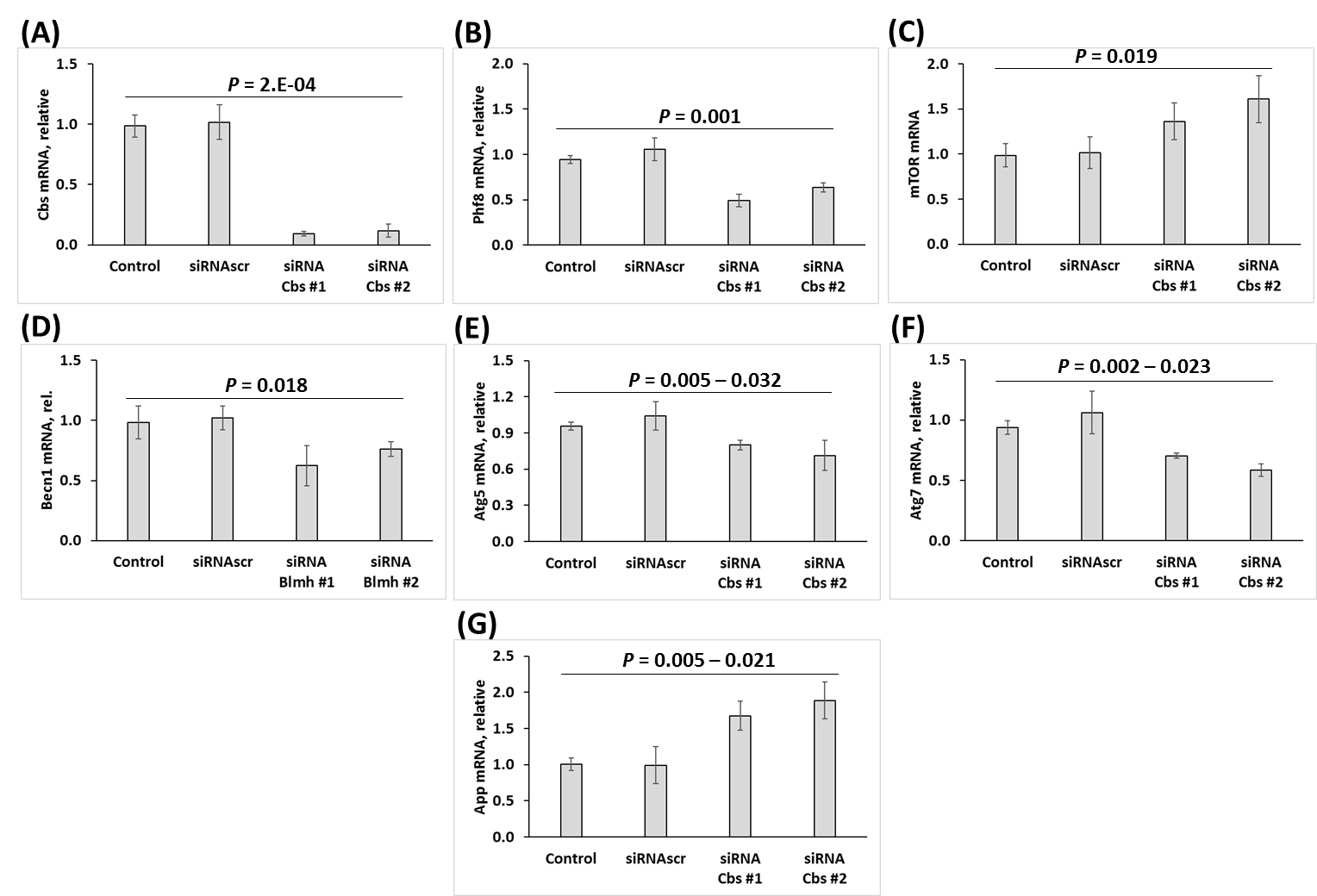

**Figure S2.** *Cbs* gene silencing affects mRNA for Phf8, mTOR, App, and autophagy-related proteins in mouse neuroblastoma N2a-APPswe cells. Bar graphs illustrating the quantification by RT-qPCR of mRNAs for Cbs (**A**), Phf8 (**B**), mTOR (**C**), Bcln1 (**D**), Atg5 (**E**), Atg7 (**F**), and App (**G**), in N2a-APPswe cells transfected with two different siRNAs targeting the *Cbs* gene (siRNA Cbs #1 and #2) are shown. Gapdh mRNA was used as a reference. Transfections without siRNA (Control) or with scrambled siRNA (siRNAscr) were used as controls.

| **Table S1 Primers used for PCR or RT-qPCR** | |
| --- | --- |
| **Gene** | **Primer sequence** |
| APP  App | Forward: 5′-CTTCCCCAAGATCCTGATAAACT-3′ |
|  | Reverse: 5′-CCGGGTGTCTCCAGGTACT-3′ |
| Atg5 | Forward: 5′-AAGGCACACCCCTGAAATGG-3′ |
|  | Reverse: 5′-TGATGTTCCAAGGAAGAGCTGAA-3′ |
| Atg7 | Forward: 5′-GCCAACTCCACACTGCTTTC-3′ |
|  | Reverse: 5′-TCTTCTGGGTCAGTTCGTGC-3′ |
| β-actin | Forward: 5′-GCAGGAGTACGATGAGTCCG-3′ |
|  | Reverse: 5′-ACGCAGCTCAGTAACAGTCC-3′ |
| Beclin-1 | Forward: 5′-GAGGAAGCTCAGTACCAGCG-3′ |
|  | Reverse: 5′-CCAGATGTGGAAGGTGGCAT-3′ |
| Cbs | Forward: 5′-GGTCTGGAATTCACTATGTAGC-3’ |
|  | Wild type reverse: 5′-CGGATGACCTGCATTCATCT-3′ Mutant reverse: 5′-GAGGTCGACGGTATCGATA-3′ |
| Gapdh | Forward: 5′-GGACTGGATAAGCAGGGCG-3′ |
|  | Reverse: 5′-TTTTGTCTACGGGACGAGGC-3′ |
| mTOR | Forward: 5′-GCCACTCTCTGACCCAGTTC 3′ |
|  | Reverse: 5′-ATGCCAAGACACAGTAGCGG-3′ |
| Phf8 | Forward: 5′-TGGGAGCATGCTTCAAGG-3′ |
|  | Reverse: 5′-GATTTCAAAGCAGGGTCATCA-3′ |
| p62 | Forward: 5′-GGGGAAGGGTTCAATGAGAG-3′ |
|  | Reverse: 5′-AATGGGCATATTTGGGGTCT-3′ |
| mTOR upstream TSS* | Forward: 5′- TTGCCAACTGGTGCTCGTTT-3′ |
|  | Reverse: 5′ AAG AAT TGG AGC TC GGG ACC 3′ |
| mTOR TSS* | Forward: 5′-GGATGTTCCTCCCCAATCTTCG-3′ |
|  | Reverse: 5′-CAGACCCACCTAACTGACCGT-3′ |
| mTOR downstream TSS* | Forward: 5′-TAGGGGGCAGATCCCGAAAC-3′ |
|  | Reverse: 5′-CACTGTAGCTGTAACTCACAC-3′ |
| * TSS, transcriprtion start site | |
